# The genome assembly and annotation of *Magnolia biondii* Pamp., a phylogenetically, economically, and medicinally important ornamental tree species

**DOI:** 10.1101/2020.06.17.158428

**Authors:** Shanshan Dong, Min Liu, Yang Liu, Fei Chen, Ting Yang, Lu Chen, Xingtan Zhang, Xing Guo, Dongming Fang, Linzhou Li, Tian Deng, Zhangxiu Yao, Xiaoan Lang, Yiqing Gong, Ernest Wu, Yaling Wang, Yamei Shen, Xun Gong, Huan Liu, Shouzhou Zhang

## Abstract

*Magnolia biondii* Pamp. (Magnoliaceae, magnoliids) is a phylogenetically, economically, and medicinally important ornamental tree species widely grown and cultivated in the north-temperate regions of China. Contributing a genome sequence for *M. biondii* will help resolve phylogenetic uncertainty of magnoliids and further understand individual trait evolution in *Magnolia*. We assembled a chromosome-level reference genome of *M. biondii* using ~67, ~175, and ~154 Gb of raw DNA sequences generated by Pacific Biosciences Single-molecule Real-time sequencing, 10X genomics Chromium, and Hi-C scaffolding strategies, respectively. The final genome assembly was ⍰2.22 Gb with a contig N50 of 269.11 Kb and a BUSCO complete gene ratio of 91.90%. About 89.17% of the genome length was organized to 19 chromosomes, resulting in a scaffold N50 of 92.86 Mb. The genome contained 48,319 protein-coding genes, accounting for 22.97% of the genome length, in contrast to 66.48% of the genome length for the repetitive elements. We confirmed a Magnoliaceae specific WGD event that might have probably occurred shortly after the split of Magnoliaceae and Annonaceae. Functional enrichment of the *Magnolia* specific and expanded gene families highlighted genes involved in biosynthesis of secondary metabolites, plant-pathogen interaction, and response to stimulus, which may improve ecological fitness and biological adaptability of the lineage. Phylogenomic analyses recovered a sister relationship of magnoliids and Chloranthaceae, which are sister to a clade comprising monocots and eudicots. The genome sequence of *M. biondii* could empower trait improvement, germplasm conservation, and evolutionary studies on rapid radiation of early angiosperms.

## Introduction

The family Magnoliaceae Juss., with over 300 species^1^ worldwide, comprises two genera, *Liriodendron* L. with only two species, and *Magnolia* L. with the rest of them^2^. About 80% of all extant Magnoliaceae species are distributed in the temperate and tropical regions of Southeast Asia, and the reminder in Americas, from temperate southeast North America through Central America to Brazil^3^, forming disjunct distribution patterns^4^.

*Magnolia* lies within magnoliids, one of the earliest assemblages of angiosperms, and occupies a pivotal position in the phylogeny of angiosperms^5^. After early divergences of angiosperms (Amborellales, Austrobaileyales, and Nymphaeales), the rapid radiation of five lineages of mesangiosperms (magnoliids, Chloranthaceae, *Ceratorpyllum*, monocots, and eudicots) occurred within a very short time frame of < 5 MYA^6^, leading to unresolved/controversial phylogenetic relationships among some lineages of mesangiosperms^5^. To date, of the 323 genome sequences available for angiosperm species^7^, mostly of plants of agronomic value, only five genomes are available for magnoliids, including black pepper^8^, avocado^9^, soursop^10^, stout camphor tree^11^, and *Liriodendron chinense*^12^. Phylogenomic analyses based on these genome data have led to controversial taxonomic placements of magnoliids. Specifically, magnoliids are resolved as the sister to eudicots with relatively strong support^11^, which is consistent with the result of the phylotranscriptomic analysis of the 92 streptophytes^13^ and of 20 representative angiosperms^14^. Alternatively, magnoliids are resolved as the sister to eudicots and monocots with weak support^8–10,12^, which is congruent with large-scale plastome phylogenomic analysis of land plants, Viridiplantae, and angiosperms^15–17^. As phylogenetic inferences rely heavily on the sampling of lineages and genes, as well as analytical methods^5^, this controversial taxonomic placements of magnoliids relative to monocots and eudicots need to be further examined with more genome data from magnoliids.

*Magnolia* species are usually cross-pollinated with precocious pistils, resulting in a very short pollination period. Many species of the genus have relatively low rates of pollen and seed germination^18^ as well as low production of fruits and seeds, which leads to difficult natural population regeneration in the wild^19–21^. Exacerbated by native habitat loss due to logging and agriculture, about 48% of all *Magnolia* species are threatened in the wild^1^. Conservation of the germplasm resources of *Magnolia*, has many economical and ecological values. Most of the *Magnolia* species are excellent ornamental tree species^22^ due to their gorgeous flowers with sweet fragrances and erect tree shape with graceful foliage, such as *M. denudata*, *M. liliiflora* and *M. grandiflora. Magnolia* species also contain a rich array of terpenoids in their flowers^23^, and have considerable varieties of phenolic compounds in their bark^24^. Many *Magnolia* species, such as *M. officinalis*, *M. biondii*, *M. denudata*, and *M. sprengeri* have been cultivated for medicinal and cosmetic purposes^25^. However, the lack of a high-quality reference genome assembly in *Magnolia* hampers current conservation and utilization efforts. The genome sequences of *Magnolia*, could greatly aid molecular breeding, germplasm conservation, and scientific research of the genus.

One *Magnolia* specie that is cultivated for ornamental, pharmaceutical, and timber purposes is *Magnolia biondii* Pamp. (Magnoliaceae, magnoliids). *M. biondii* is a deciduous tree species widely grown and cultivated in the north-temperate regions of China. Its flowers are showy and fragrant and can be used to extract essential oils. The chemical extracts of the flower buds are used for local stimulation and anesthesia, anti-inflammatory, antimicrobial, analgesic, blood pressure-decreasing, and anti-allergic effects^25^. Modern phytochemical studies have characterized the chemical constitutes of the volatile oil^26^, lignans^27^, and alkaloids^28^ from different parts of the *M. biondii* plant. The volatile oils contain a rich array of terpenoids, among which, the main ingredients are 1,8-cineole, β-pinene, α-terpineol, and camphor^25^. These terpenoids are synthesized by the terpene synthase (TPS) that belongs to the TPS gene family. In this study, we sequenced and assembled the reference genome of *M. biondii* using the Pacbio long reads, 10X Genomics Chromium, and Hi-C scaffolding strategies. The ~2.22 Gb genome sequence of *M. biondii* represented the largest genome assembled to date in the early-diverging magnoliids. This genome will support future studies on floral evolution and biosynthesis of the primary and secondary metabolites unique to the species, and will be an essential resource for understanding rapid changes that took place at the backbone phylogeny of the angiosperms. Finally, it could further genome-assisted improvement for cultivation and conservation efforts of *Magnolia*.

## Materials and Methods

### Plant materials, DNA extractions, and sequencing

Fresh leaves and flower materials from three development stages were collected from a 21-year old *M. biondii* tree (a cultivated variety) planted in the Xi’an Botanical Garden, Xi’an, China. The specimen (voucher number: Zhang 201801M) has been deposited in the Herbarium of Fairy Lake Botanical Garden, Shenzhen, China. Total genomic DNA was extracted from fresh young leaves of *M. biondii* using modified cetyltrimethylammonium bromide (CTAB) method^29^. The quality and quantity of the DNA samples were evaluated using a NanoDrop™ One UV-Vis spectrophotometer (Thermo Fisher Scientific, USA) and a Qubit^®^ 3.0 Fluorometer (Thermo Fisher Scientific, USA). Three different approaches were used in genomic DNA sequencing at BGI-Shenzhen (BGI Co. Ltd., Shenzhen, China) (**Supplementary Table S1**). First, high molecular weight genomic DNA was prepared for 10X Genomics libraries with insert sizes of 350–500 bp according to the manufacturer’s protocol (Chromium Genome Chip Kit v1, PN-120229, 10X Genomics, Pleasanton, USA). The barcoded library was sequenced on a BGISEQ-500 platform to generate 150-bp read pairs. Duplicated reads, reads with ≥20% low-quality bases or with ≥5% ambiguous bases (“N”) were filtered using SOAPnuke v.1.5.6^30^ with the parameters “-l 10 -q 0.1 -n 0.01 -Q 2 -d --misMatch 1 --matchRatio 0.4 -t 30,20,30,20”. Second, single-molecule real-time (SMRT) Pacific Biosciences (PacBio) libraries were constructed using the PacBio 20-kb protocol (https://www.pacb.com/) and sequenced on a PacBio RS-II instrument. Third, a Hi-C library was generated using DpnII restriction enzyme following in situ ligation protocols^31^. The DpnII-digested chromatin was end-labeled with biotin-14-dATP (Thermo Fisher Scientific, Waltham, MA, USA) and used for in situ DNA ligation. The DNA was extracted, purified, and then sheared using Covaris S2 (Covaris, Woburn, MA, USA). After A-tailing, pull-down, and adapter ligation, the DNA library was sequenced on a BGISEQ-500 to generate 100-bp read pairs.

### RNA extraction and sequencing

Young leaves (LEAF), opening flowers (FLOWER), and flower buds (BUDA and BUDB) from two developmental stages (pre-meiosis and post-meiosis) were collected from the same individual tree planted in Xi’an Botanical Garden. Total RNAs were extracted using E.Z.N.A.^®^ Total RNA Kit I (Omega Bio-Tek) and then quality controlled using a NanoDrop™ One UV-Vis spectrophotometer (Thermo Fisher Scientific, USA) and a Qubit® 3.0 Fluorometer (Thermo Fisher Scientific, USA). All RNA samples with integrity values close to 10 were selected for cDNA library construction and next generation sequencing. The cDNA library was prepared using the TruSeq RNA Sample Preparation kit v2 (Illumina, San Diego, CA, USA) and paired-end (150 bp) sequenced on a HiSeq 2000 platform (Illumina Inc, CA, USA) at Majorbio (Majorbio Co. Ltd., Shanghai, China). The newly generated raw sequence reads were trimmed and filtered for adaptors, low quality reads, undersized inserts, and duplicated reads using Trimmomatic v. 0.38^32^.

### Genome size estimation

We used 17 k-mer counts^33^ of high-quality reads from small insert libraries of 10X genomics to evaluate the genome size and the level of heterozygosity. First, K-mer frequency distribution analyses were performed following Chang *et al.*^34^ to count the occurrence of k-mers based on the clean paired-end 10X genomics data. Then, GCE^35^ was used to estimate the general characteristics of the genome, including total genome size, repeat proportions, and level of heterozygosity (**Supplementary Table S2**).

### *De novo* genome assembly and chromosome construction

*De novo* assembly was performed with five different genome assemblers, Canu v. 0.1^36^, Miniasm v. 0.3^37^, Wtdbg v. 1.1.006 (https://github.com/ruanjue/wtdbg), Flye v. 2.3.3^38^, and SMARTdenovo 1.0.0 (https://github.com/ruanjue/smartdenovo) with/without priori Canu correction with default parameters. Based on the size of the assembled genome, the total number of assembled contigs, the length of contig N50, maximum length of the contigs, and also the completeness of the genome assembly as assessed by using Benchmarking Universal Single-Copy Orthologs (BUSCO) analysis^39^ (1,375 single copy orthologs of the Embryophyta odb10 database) with the BLAST e-value cutoff of 1e–5, genome assembly from Miniasm assembler was selected for further polishing and scaffolding (**Supplementary Table S3**). The consensus sequences of the assembly were further improved using all the PacBio reads for three rounds of iterative error correction using software Racon v. 1.2.1^40^ with the default parameters and the resultant consensus sequences were further polished using Pilon v. 1.22^41^ (parameters: --fix bases, amb --vcf --threads 32) with one round of error correction using all the clean paired-end 10X genomics reads. Hi-C reads were quality-controlled (**Supplementary Table S2**) and mapped to the contig assembly of *M. biondii* using Juicer^42^ with default parameters. Then a candidate chromosome-length assembly was generated automatically using the 3D-DNA pipeline^43^ (parameters: −m haploid −s 4 −c 19 −j 15) to correct mis-joins, order, orient, and organize contigs from the draft chromosome assembly. Manual check and refinement of the draft assembly was carried out in Juicebox Assembly Tools^44^ (**Table 1**).

**Table 1.** Final genome assembly based on the assembled contigs from Miniasm.

|  | PacBio<br>Assembly<br>(polished) | Hi-C<br>Assembly |
| --- | --- | --- |
| Total scaffold length (Gb) |  | 2.232 |
| Number of scaffolds |  | 9510 |
| Scaffold N50 (Mb) |  | 92.86 |
| Scaffold N90 (Mb) |  | 19.29 |
| Max scaffold len(Mb) |  | 168.50 |
| Total Contig length (Gb) | 2.22 |  |
| Number of contigs | 15,615 |  |
| Contig N50 (Kb) | 269.114 |  |
| Contig N90 (Kb) | 60.09 |  |
| Max contig len(Kb) | 2,134.98 |  |
| Complete BUSCOs | 91.90% | 88.50% |
| Complete and single-copy<br>BUSCOs | 87.00% | 85.20% |
| Complete and duplicated<br>BUSCOs | 4.90% | 3.30% |
| Fragmented BUSCOs | 3.00% | 4.40% |

### Genome evaluation

The completeness of the genome assembly of *M. biondii* was evaluated with DNA and RNA mapping results, transcript unigene mapping results, and BUSCO analysis^39^. First, all the paired-end reads from 10X genomics and Hi-C were mapped against the final assembly of *M. biondii* using BWA-MEM v. 0.7.10^45^. The RNA-seq reads from four different tissues were also mapped back to the genome assembly using TopHat v. 2.1.0^46^. Second, unigenes were generated from the transcript data of *M. biondii* using Bridger software^47^ with the parameters “–kmer length 25 – min kmer coverage 2” and then aligned to the scaffold assembly using Basic Local Alignment Search Tool (BLAST)- like alignment tool BLAT^48^. Third, BUSCO analysis^39^ of the final scaffold assembly were also performed to evaluate the genome completeness of the reference genome of *M. biondii*.

### Repeat annotation

Transposable elements (TEs) were identified by a combination of homology-based and *de novo* approaches. Briefly, the genome assembly was aligned to a known repeats database Repbase v. 21.01^49^ using RepeatMasker v. 4.0.5^50^ and Repeat-ProteinMask^50^ at both the DNA and protein level for homology-based TE characterization. In the *de novo* approach, RepeatModeler 2.0^51^ and LTR Finder v. 1.0.6^52^ were used to build a *de novo* repeat library using the *M. biondii* assembly. TEs in the genome were then identified and annotated by RepeatMasker v. 4.0.5^50^. Tandem repeats were annotated in the genome using TRF v. 4.04^53^ (**Supplementary Table S4**).

### Gene prediction

Protein-coding genes were predicted by using the MAKER-P pipeline v. 2.31^54^ based on *de novo* prediction, homology search, and transcriptome evidences. For *de novo* gene prediction, GeneMark-ES v. 4.32^55^ was firstly used for self-training with the default parameters. Secondly, the alternative spliced transcripts, obtained by a genome-guided approach by using Trinity with the parameters “--full_cleanup --jaccard_clip --no_version_check --genome_guided_max_intron 100000 --min_contig_length 200” were mapped to the genome by using PASA v. 2.3.3 with default parameters. Then the complete gene models were selected and used for training Augustus^56^, and SNAP^57^. They were used to predict coding genes on the repeat-masked *M. biondii* genome. For homologous comparison, protein sequences from *Arabidopsis thaliana*, *Oryza sativa*, *Amborella trichopoda*, and two related species (*C. kanehirae* and *L. chinense*) were provided as protein evidences.

For RNA evidence, a completely *de novo* approach was chosen. The clean RNA-seq reads were then assembled into inchworm contigs using Trinity v. 2.0.6^58^ with the parameters “--min_contig_length 100 --min_kmer_cov 2 --inchworm_cpu 10 --bfly_opts “-V 5 --edge-thr=0.05 --stderr” --group_pairs_distance 200 --no_run_chrysalis” and then provided to MAKER-P as expressed sequence tag evidence. After two rounds of MAKER-P, a consensus gene set was obtained. tRNAs were identified using tRNAscan-SE v. 1.3.1^59^. snRNA and miRNA were detected by searching the reference sequence against the Rfam database^60^ using BLAST^61^. rRNAs were detected by aligning with BLASTN^61^ against known plant rRNA sequences^62^ (**Supplementary Table S5**). We also mapped the gene density, GC content, *Gypsy* density, and *Copia* density on the individual chromosomes using Circos tool (http://www.circos.ca) (**Fig. 1**).

**Fig. 1.**
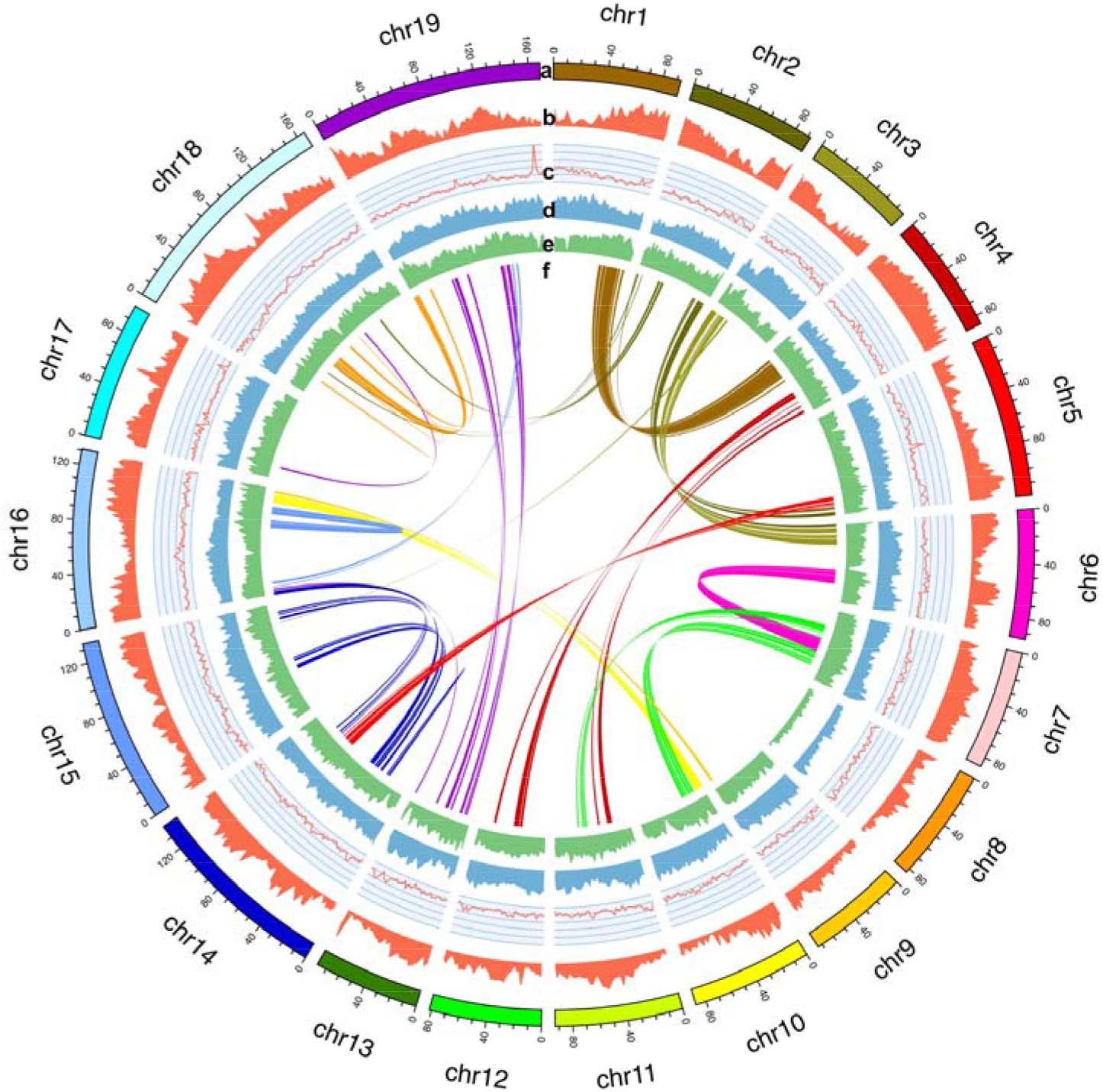
Reference genome assembly of nighteen chromosomes. **a.** Assembled chromosomes, **b.** Gene density, **c.** GC content, **d.** *Gypsy* density, **e.** *Copia* density, and **f.** Chromosome synteny (from outside to inside).

### Functional annotation of protein-coding genes

Functional annotation of protein-coding genes was performed by searching the predicted amino acid sequences of *M. biondii* against the public databases based on sequence identity and domain conservation. Protein-coding genes were previously searched against the following protein sequence databases, including the Kyoto Encyclopedia of Genes and Genomes (KEGG)^63^, the National Center for Biotechnology Information (NCBI) non-redundant (NR) and the Clusters of Orthologous Groups (COGs) databases^64^, SwissProt^65^, and TrEMBL^65^, for best matches using BLASTP with an e-value cutoff of 1e−5. Then, InterProScan 5.0^66^ was used to characterize protein domains and motifs based on Pfam^67^, SMART^68^, PANTHER^69^, PRINTS^70^, and ProDom^71^ (**Supplementary Table S6**).

### Gene family construction

Protein and nucleotide sequences from *M. biondii* and six other angiosperms plants (*Amborella trichopoda*, *Arabidopsis thaliana*, *Cinnamomum Kanehirae*, *Liriodendron chinense*, *Vitis vinifera*) were used to construct gene families using OrthoFinder^72^ (https://github.com/davidemms/OrthoFinder) based on an all-versus-all BLASTP alignment with an e-value cutoff of 1e−5. Potential gene pathways were obtained via gene mapping against the KEGG databases, and Gene Ontology (GO) terms were extracted from the corresponding InterProScan or Pfam results (**Fig. 2**).

**Fig. 2.**
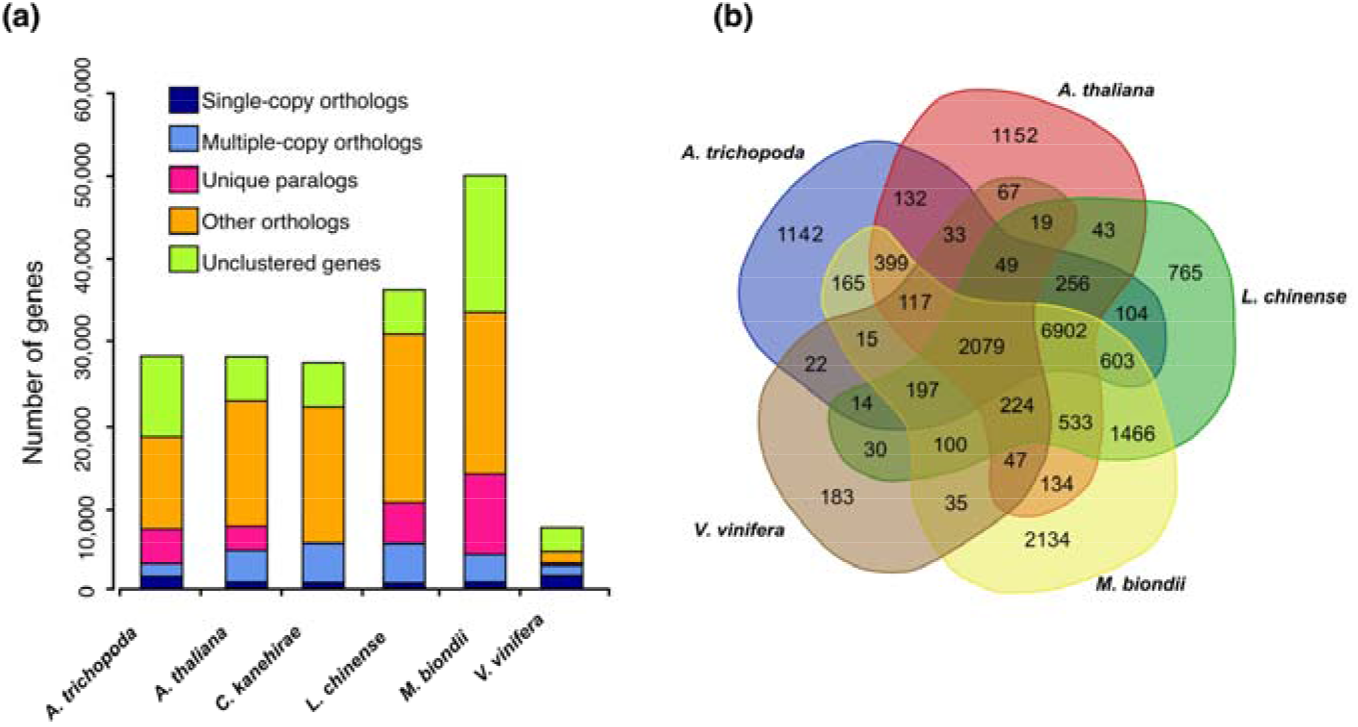
Comparative analysis of the *M. biondii* genome. **(a)** The number of genes in various plant species, showing the high gene number of *M. biondii* compared to a model (*Arabidopsis thaliana*) and other species (including *Amborella trichopoda*, *Cinnamomum kanehirae*, *Liriodendron chinense*, and *Vitis vinifera*). **(b)** Venn diagram showing overlaps of gene families between *M. biondii*, *L. chinense*, *A. trichopoda*, *A. thaliana*, and *V. vinifera*.

### Phylogenomic reconstruction and gene family evolution

To understand the relationships of the *M. biondii* gene families with those of other plants and the phylogenetic placements of magnoliids among angiosperms, we performed a phylogenetic comparison of genes among different species along a 20-seed plant phylogeny reconstructed with a concatenated amino acid dataset derived from 109 single-copy nuclear genes. Putative orthologous genes were constructed from 18 angiosperms (including two eudicots, two monocots, two Chloranthaceae species, eight magnoliid species, two *Illicium* species, *A. trichopoda*, *Nymphaea sp.*) and the gymnosperm outgroup *Picea abies* (**Supplementary Table S7**) using OrthoFinder^72^ and compared with protein genes from the genome assembly of *M. biondii*. The total of one-to-one orthologous gene sets were identified and extracted for alignment using Mafft v. 5.0^73^, further trimmed using Gblocks 0.91b^74^, and concatenated in Geneious 10.0.2 (www.geneious.com). The concatenated amino acid dataset from 109 single copy nuclear genes (each with >85% of taxon occurrences) was analyzed using PartitionFinder^75^ with an initial partitioning strategy by each gene for optimal data partitioning scheme and associated substitution models, resulting in 18 partitions. The concatenated amino acid dataset was then analyzed using the maximum likelihood (ML) method with RAxML-VI-HPC v. 2.2.0^76^ to determine the best reasonable tree. Non-parametric bootstrap analyses were implemented by PROTGAMMALG approximation for 500 pseudoreplicates (**Fig. 3**).

**Fig. 3.**
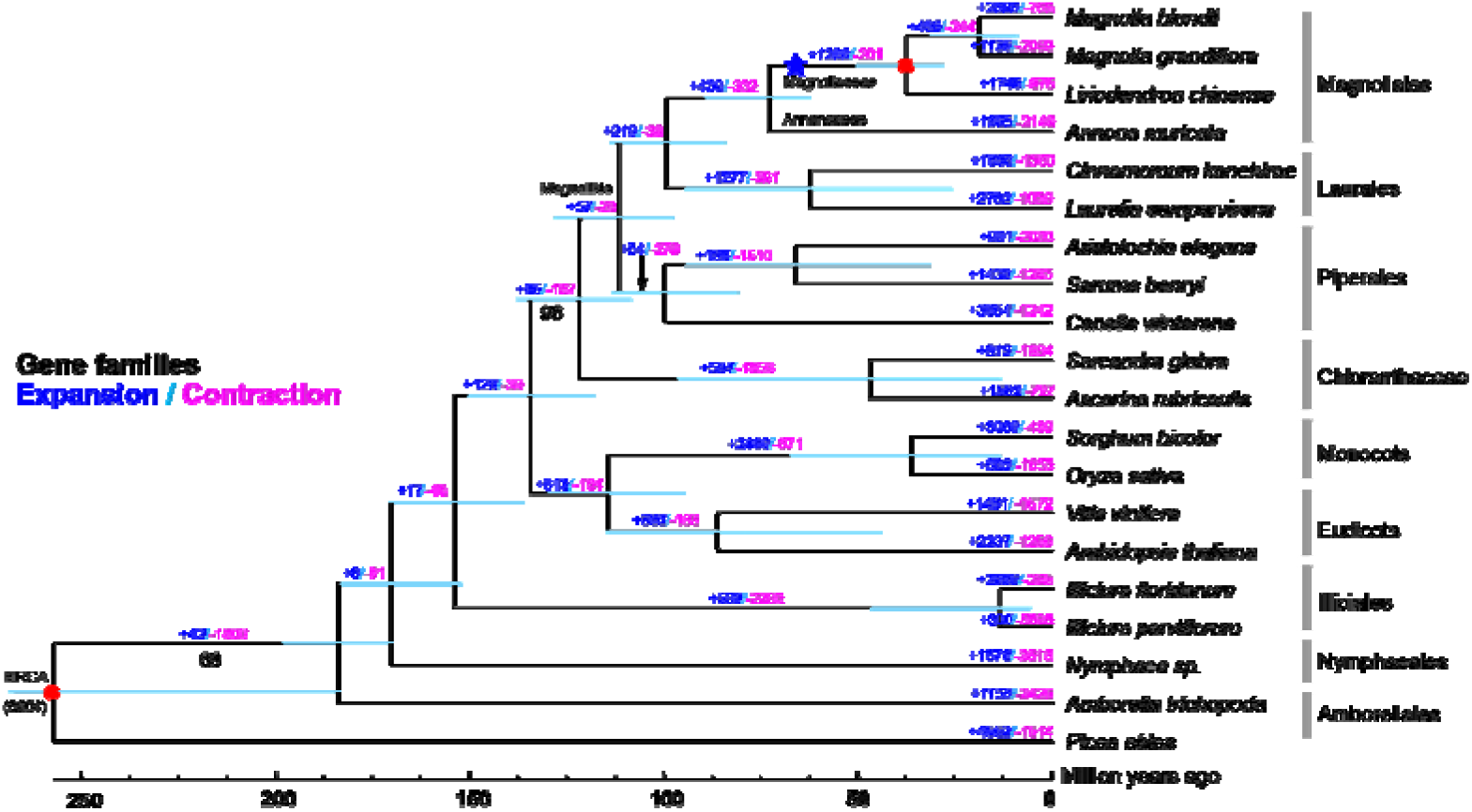
Phylogenetic tree and number of gene families displaying expansions and contractions among 20 plant species. Estimated divergence time confidence intervals are shown at each internal node as teal bars. Calibrated nodes are indicated by red dots. The Magnoliaceae specific WGD is indicated with blue stars. All the branches are maximally supported by maximum likelihood analysis unless otherwise indicated below the branches.

The best maximum likelihood tree was used as a starting tree to estimate species divergence time using MCMC Tree as implemented in PAML v. 4^77^. Two node calibrations were defined from the Timetree web service (http://www.timetree.org/), including the split between *Liriodendron* and *Magnolia* (34–77 MYA) and the split between angiosperms and gymnosperms (168–194 MYA). The orthologous gene clusters inferred from the OrthoFinder^72^ analysis and phylogenetic tree topology constructed using RAxML-VI-HPC v. 2.2.0^76^ were taken into CAFE v. 4.2^78^ to indicate whether significant expansion or contraction occurred in each gene family across species.

### Analyses of genome synteny and whole-genome duplication (WGD)

To investigate the source of the large number of predicted protein genes (48,319) in *M. biondii*, the whole genome duplication (WGD) events were analyzed by making use of the high-quality genome of *M. biondii*. As the grape genome have one well-established whole-genome triplication, and the co-familial *L. chinense* have one reported whole genome duplication event^12^, the protein-coding genes (of CDS and the translated protein sequences, respectively) of *M. biondii* with that of itself, *L. chinense*, and the grape were used to perform synteny searches with MCscanX^79^(python version), with at least five gene pairs required per syntenic block. The resultant dot plots were examined to predict the paleoploidy level of *M. biondii* in comparison to the other angiosperm genomes by counting the syntenic depth in each genomic region (**Supplementary Fig. S3, S4**). The synonymous substitution rate (Ks) distribution for paralogues found in collinear regions (anchor pairs) of *M. biondii* and *chinense*, was analyzed with WGD suite^80^ with default parameters (**Fig. 4**).

**Fig. 4.**
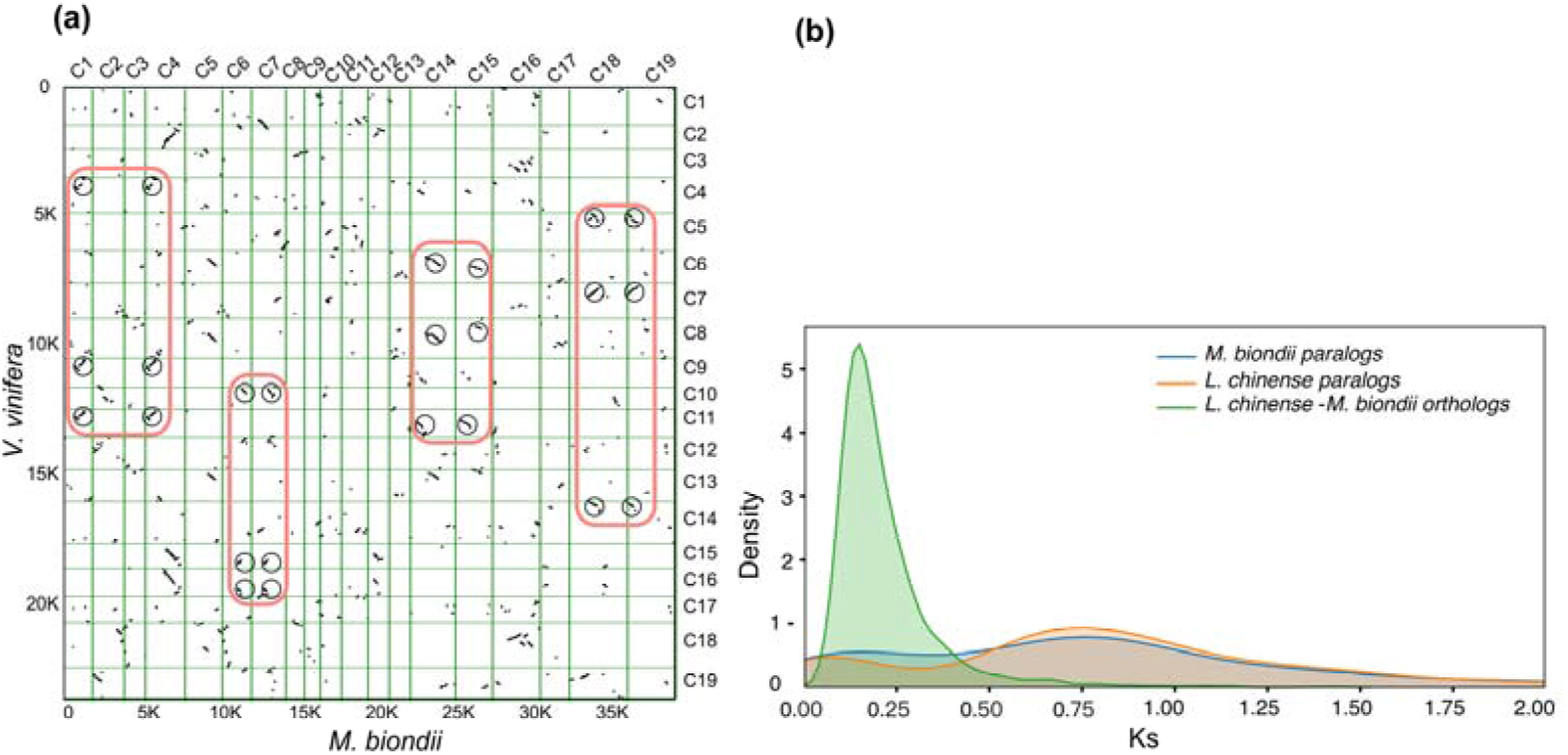
Evidences for whole-genome duplication events in *M. biondii*. **(a)** Comparison of *M. biondii* and grape genomes. Dot plots of orthologues show a 2–3 chromosomal relationship between the *M. biondii* genome and grape genome. **(b)** Synonymous substitution rate (Ks) distributions for paralogues found in collinear regions (anchor pairs) of *M. biondii* and *Liriodendron chinense*, and for orthologues between *M. biondii* and *L. chinense*, respectively.

### Identification of TPS genes and Expression analysis

We selected two species (*A. trichopoda*, *A. thaliana*) to perform comparative TPS gene family analysis with *M. biondii*. Previously annotated TPS genes of two species were retrieved from the data deposition of Chaw *et al.*^11^. Two Pfam domains: PF03936 and PF01397, were used as queries to search against the *M. biondii* proteome using HMMER v. 3.0 with an e-value cut-off of 1e−5^82^. Protein sequences with lengths below 200 amino acids were removed from subsequent phylogenetic analysis. Putative protein sequences of TPS genes were aligned using MAFFT v. 5^73^ and manually adjusted using MEGA v. 4^83^. The phylogenetic tree was constructed using IQ-TREE^84^ with 1,000 bootstrap replicates (**Fig. 5**).

**Fig. 5.**
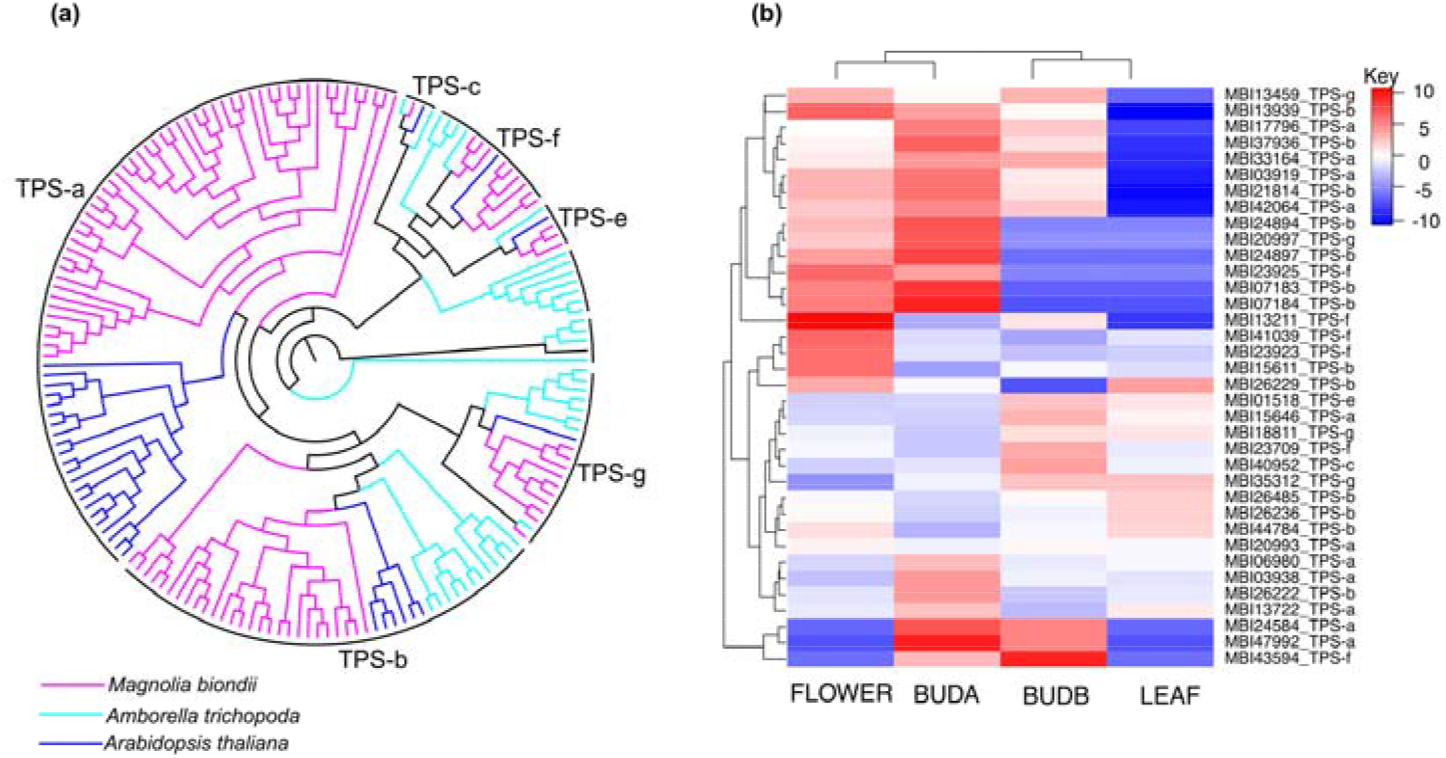
TPS (terpene synthase) gene family in *M. biondii*. (**a**) The phylogenetic tree of TPS genes from *Amborella trichopoda*, *Arabidopsis thaliana*, and *M. biondii*. (**b)** Heatmap showing differential expression of TPS genes in the transcriptome data from young leaves (LEAF), opening flowers (FLOWER), flower buds of pre-meiosis (BUDA), and flower buds of post-meiosis (BUDB).

### Data access

The genome assembly, annotations, and other supporting data are available at dryad database under the DOI: https://doi.org/10.5061/dryad.s4mw6m947. The raw sequence data have been deposited in the China National GeneBank DataBase (CNGBdb) under the Accession No. of CNP0000884.

## Results

### Sequencing summary

DNA sequencing generated 33-fold PacBio single-molecule long reads (a total of 66.78 Gb with an average length of 10.32 kb), 80-fold 10X genomics paired-end short reads (175.45 Gb) and Hi-C data (~153.78 Gb). Transcriptome sequencing generated 4.62, 4.60, 4.67, and 4.73 Gb raw data for young leaves, opening flowers, and flower buds from two developmental stages (pre-meiosis and post-meiosis), respectively (**Supplementary Table S1**).

### Determination of genome size and heterozygosity

K-mer frequency distribution analyses suggested a k-mer peak at a depth of 48, and an estimated genome size of 2.17 Gb (**Supplementary Fig. S1a**, **Table S2**). GCE^35^ analysis resulted in a k-mer peak at a depth of 29, and a calculated genome size of 2.24 Gb, an estimated heterozygosity of 0.73%, and a repeat content of 61.83% (**Supplementary Fig. S1b**, **Table S2**). The estimated genome size of *M. biondii* is the largest among all the sequenced genomes of magnoliids.

### Genome assembly and quality assessment

The selected primary assembly from Miniasm v. 0.3^37^ has a genome size of 2.20 Gb across 15,713 contigs, with a contig N50 of 267.11 Kb. After three rounds of error correction with Pacbio long reads and one round of correction with 10X genomics reads, we arrived at a draft contig assembly size of 2.22 Gb spanning 15,628 contigs with a contig N50 of 269.11 Kb (**Table 1**). About 89.17% of the contig length was organized to the 19 chromosomes (1.98 Gb), with ambiguous Ns accumulated to 7,365,981 bp (accounting for 0.33% of the genome length). About 9,455 contigs (0.24 Gb) were unplaced (**Supplementary Fig. S2**). The raw scaffold assembly was further improved with Pacbio long reads and 10X genomics short reads, resulting in an assembled genome size of 2.23 Gb represented by 9,510 scaffolds with a scaffold N50 of 92.86 Mb (**Table 1**). Our assembled genome size of *M. biondii* is very much approximate to the estimated genome size of K-mer analysis (**Supplementary Table S2**).

For genome quality assessment, First, all the paired-end reads from 10X genomics and Hi-C were mapped against the final assembly of *M. biondii*, resulting in 98.40% and 92.50% of the total mapped reads, respectively. Sequencing coverage of 10X genomics reads and Hi-C reads showed that more than 98.04% and 86.00% of the genome bases had a sequencing depth of >10×, respectively. The RNA-seq reads from four different tissues were also mapped back to the genome assembly using TopHat v. 2.1.0^46^, resulting in 93.3%, 94.4%, 92.9%, and 93.7% of the total mapped RNA-seq reads for leaves, opening flowers, flower buds of pre-meiosis and post-meiosis, respectively. Second, unigenes generated from the transcript data of *M. biondii* were aligned to the scaffold assembly. The result indicated that the assemblies covered about 86.88% of the expressed unigenes. Third, BUSCO analysis^39^ of the final scaffold assembly showed that 88.50% (85.20% complete and single-copy genes and 3.30% complete and duplicated genes) and 4.40% of the expected 1,375 conserved embryophytic genes were identified as complete and fragmented genes, respectively. These DNA/RNA reads and transcriptome unigene mapping studies, and BUSCO analysis suggested an acceptable genome completeness of the reference genome of *M. biondii*.

### Repeat annotation

We identified 1,478,819,185 bp (66.48% of the genome length) bases of repetitive sequences in the genome assemblies of *M. biondii*. LTR elements were the predominant repeat type, accounting for 58.06% of the genome length (**Supplementary Table S4**). For the two LTR superfamilies, *Copia* and *Gypsy* elements accumulated to 659,463,750 and 727,531,048 bp, corresponding to 45.26% and 50.66% of the total LTR repeat length, respectively. The density of *Gypsy* elements scaled negatively with the density of genes whereas *Copia* elements distributed more evenly across the genome and showed no obvious patterns or correlationships with the distribution of genes (**Fig. 1**). DNA transposons, satellites, simple repeats and other repeats accumulated to 130,503,028, 5,540,573, 17,626,796, and 7,240,517 bp, accounting for 5.86%, 0.24%, 0.79%, and 0.32% of the genome length, respectively.

### Gene annotation and functional annotation

The assembled genome of *M. biondii* contained 48,319 protein-coding genes, 109 miRNAs, 904 tRNAs, 1,918 rRNAs, and 7,426 snRNAs (**Supplementary Table S5**). The protein-coding genes in *M. biondii* had an average gene length of 10,576 bp, an average coding DNA sequence (CDS) length of 950 bp, and an average exon number per gene of 4.4. Various gene structure parameters were compared to those of the five selected species, including *A. trichopoda*, *A. thaliana*, *C. kanehirae*, *L. chinense*, and *Oryza sativa. M. biondii* had the highest predicted gene numbers and the largest average intron length (~2,797 bp) among these species (**Supplementary Table S5**), which appears to be in agreement with the relatively larger genome size of *M. biondii*. However, the relatively small median gene length (3,390 bp) and intron length (532 bp) in *M. biondii* suggested that some genes with exceptionally long introns have significantly increased the average gene length.

Functional annotation of protein-coding genes assigned potential functions to 39,405 protein-coding genes out of the total of 48,319 genes in the *M. biondii* genome (81.55 %) (**Supplementary Table S6**). Among ~18.5% of the predicted genes without predicted functional annotations, some may stem from errors in genome assembly and annotations, while others might be potential candidates for novel functions.

### Gene family construction

Among a total of 15,150 gene families identified in the genome of *M. biondii*, 10,783 genes and 1,983 gene families were found specific to *M. biondii* (**Fig. 2a**). The Venn diagram in **Fig. 2b** shows that 2,079 gene families were shared by the five species, *M. biondii*, *L. chinense*, *A. trichopoda*, *A. thaliana*, and *V. vinifera*. Specific gene families were also detected in these five species. A total of 11,057 genes and 2,134 gene families were found to be specific to *M. biondii*.

A KEGG pathway analysis of the *M. biondii* specific gene families revealed marked enrichment in genes involved in nucleotide metabolism, plant-pathogen interaction, and biosynthesis of alkaloid, ubiquinone, terpenoid-quinone, phenylpropanoid, and other secondary metabolites (**Supplementary Table S8**), which is consistent with the biological features of *M. biondii* with rich arrays of terpenoids, phenolics, and alkaloids. Using Gene Ontology (GO) analysis, the *M. biondii* specific gene families are enriched in binding, nucleic acid binding, organic cyclic compound binding, heterocyclic compound binding, and hydrolase activity (**Supplementary Table S9**). The specific presence of these genes associated with biosynthesis of secondary metabolites and plant-pathogen interaction in *M. biondii* genome assembly might play important roles in plant pathogen-resistance mechanisms^8^ by stimulating beneficial interactions with other organisms^11^.

### Phylogenomic reconstruction

Our phylogenetic analyses based on 109 orthologous nuclear single-copy genes and 19 angiosperms plus one gymnosperm outgroup recovered a robust topology and supported the sister relationship of magnoliids and Chloranthaceae (BPP=96), which together formed a sister group relationship (BPP=100) with a clade comprising monocots and eudicots. The phylogenetic tree (**Fig. 3**) indicates that the orders of Magnoliales and Laurales have a close genetic relationship, with a divergence time of ~99.3 MYA (84.4–115.5 MYA). The estimated divergence time of Magnoliaceae and Annonaceae in the Magnoliales clade is ⍰72.8 MYA (56.5–91.5 MYA), while the split of *Liriodendron* and *Magnolia* is estimated at ~37.6 MYA (31.3–50.2 MYA).

### Gene family evolution

The orthologous gene clusters inferred from the OrthoFinder^72^ analysis and phylogenetic tree topology constructed using RAxML-VI-HPC v. 2.2.0^76^ were taken into CAFE v. 4.2^78^ to indicate whether significant expansion or contraction occurred in each gene family across species (**Fig. 3**). Among a total of 15,683 gene families detected in the *M. biondii* genome, 2,395 were significantly expanded (P < 0.05) and 765 contracted (P<0.005). A KEGG pathway analysis of these expanded gene families revealed marked enrichment in genes involved in metabolic pathways, biosynthesis of secondary metabolites, plant hormone signal transduction, ABC transporters and etc. (**Supplementary Table S10**). Using Gene Ontology (GO) analysis, the *M. biondii* expanded gene families are enriched in ion binding, transferase activity, metabolic process, cellular process, oxidoreductase activity, localization, response to stimulus, and etc. (**Supplementary Table S11**). The expansion of these genes especially those associated with biosynthesis of secondary metabolites, plant hormone signal transduction and response to stimulus might possibly contribute to the ecological fitness and biological adaptability of the species.

### Analyses of genome synteny and whole-genome duplication (WGD)

A total of 1,715 colinear gene pairs on 144 colinear blocks were inferred within the *M. biondii* genome (**Supplementary Fig. S4a**). There were 13,630 co-linear gene pairs from 393 colinear blocks detected between *M. biondii* and *L. chinense* (**Supplementary Fig. S4b**), and 9,923 co-linear gene pairs from 915 co-linear blocks detected between *M. biondii* and *V. vinifera* (**Fig. 4a**). Dot plots of longer syntenic blocks between *M. biondii* and *L. chinense* revealed a nearly 1:1 orthology ratio, indicating a similar evolutionary history of *M. biondii* to *L. chinense. Magnolia* may probably have also experienced a WGD event as *Liriodendron*^12^ after the most recent common ancestor (MRCA) of angiosperms. And that, the nearly 2:3 orthology ratio between *M. biondii* and grape confirmed this WGD event in the lineage leading to *Magnolia* (**Supplementary Fig. S4b**).

The Ks distribution for *M. biondii* paralogues revealed a main peak at around 0.75 Ks (~124 Ma) units, which appears to coincide with the Ks peak of *L. chinense* in our observation (**Fig. 4b**), indicating that the two lineages might have experienced a shared WGD in their common ancestor or two independent WGDs at a similar time. The one-vs-one ortholog comparisons between *Liriodendron* and *Magnolia* suggested the divergence of the two lineages at around 0.18 Ks units, which largely postdates the potential WGD peak of 0.75 Ks units observed in either species, indicating that this WGD event should be shared at least by the two genera of Magnoliaceae.

### TPS genes

The volatile oils isolated from the flower buds of *M. biondii* constitute primarily terpenoid compounds that are catalyzed by TPS enzymes. We identified a total of 102 putative TPS genes in the genome assembly of *M. biondii*, which is comparable to that of the *C. kanehirae* with 101 genes^11^. To determine the classification of TPS proteins in *M. biondii*, we constructed a phylogenetic tree using all the TPS protein sequences from *M. biondii*, *A. thaliana* and *A. trichopoda*. These TPS genes found in *biondii* can be assigned to six subfamilies, TPS-a (52 genes), TPS-b (27 genes), TPS-c (1 gene), TPS-e (3 genes), TPS-g (3 genes), and TPS-f (9 genes) (**Fig. 5a**). We compared the expression profiles of TPS genes in the young leaves and flowers from three different developmental stages (**Fig. 5b**), and identified a total of 36 TPS genes (including 11, 13, 1, 1, 6, and 4 genes for the subfamilies of TPS-a, TPS-b, TPS-c, TPS-e, TPS-f, and TPS-g, respectively) substantially expressed, among which, 33 TPS genes (including both 10 genes for TPS-a and TPS-b subfamilies) exhibited higher transcript abundance in flowers, compared to leaves (**Fig. 5b**), suggesting that these genes may be involved in a variety of terpenoid metabolic processes during flower growth and development in *M. biondii*.

## Discussion

The genome of *M. biondii* is relatively large and complex as K-mer frequency analysis suggested an estimated genome size of 2.24 Gb, with an estimated heterozygosity of 0.73%, and a repeat content of 61.83%. Our DNA sequencing generated about 33-fold PacBio long reads data, which resulted in an assembly of 2.23 Gb spanning 15,628 contigs with a contig N50 of 269.11 Kb. The small contig N50 length might imply fragmentary and incomplete genome assembly, which might affect the quality and precision of the Hi-C assembly. Indeed, when these contigs were organized to chromosomes using Hi-C data, about 6,899 contigs adding up to 1.00 Gb were disrupted by the Hi-C scaffolding processes, contributing to 0.18 Gb genome sequences discarded. After manual correction of the Hi-C map in Juicer box, the final scaffold assembly has still 6,911 contigs disrupted, 2,358 genes disturbed, and 0.24 Gb of genome sequences unplaced. BUSCO assessments show decreased percentages of complete BUSCOs and increased percentages of fragmented BUSCOs for the scaffold assembly than that of the contig assembly (**Table 1**). Therefore, we used the HiC assembly for chromosome collinearity analysis and the contig assembly for the rest of comparative analyses. The exceptionally large protein gene set predicted for *M. biondii* genome might be attributed to gene fragmentation problems induced by poor genome assembly and high content of transposable elements, as evidenced by dramatically short average/median CDS length of *M. biondii* compared with that of the co-familial *L. chinense* (Supplementary Table S5).

The chromosome-scale reference genome of *M. biondii* provided information on the gene contents, repetitive elements, and genome structure of the DNA in the 19 chromosomes. Our genome data offered valuable genetic resources for molecular and applied research on *M. biondii* as well as paved the way for studies on evolution and comparative genomics of *Magnolia* and the related species. Phylogenomic analyses of 109 single-copy orthologues from 20 representative seed plant genomes with a good representation of magnoliids (three out of four orders) strongly support the sister relationship of magnoliids and Chloranthaceae, which together form a sister group relationship with a clade comprising monocots and eudicots. This placement is congruent with the plastid topology^15,16^ and the multi-locus phylogenetic studies of angiosperms^6^, but in contrast to the placement of the sister group relationship of magnoliids with eudicots recovered by the phylogenomic analysis of angiosperms (with *Cinnamomum kanehirae* as the only representative for magnoliids)^11^, phylotranscriptomic analysis of the 92 streptophytes^13^ and of 20 representative angiosperms^14^. Multiple factors underlies the robust angiosperm phylogeny recovered in our study: (a) we use less homoplasious amino acid data rather than nucleotide sequences (especially those of the 3^rd^ codon positions) that are more prone to substitution saturation; (b) we use an optimal partitioning strategy with carefully selected substitution models, which is usually neglected for large concatenated datasets in phylogenomic analyses; (c) we have a relatively good taxa sampling that included representatives from all the eight major angiosperm lineages but Ceratophyllales that has no genome resources available. Future phylogenomic studies with an improved and more balanced lineage sampling and a thorough gene sampling as well as comprehensive analytical methods would provide more convincing evidences on the divergence order of early mesangiosperms.

The current assembly of the *M. biondii* genome informed our understanding of the timing of the WGD event in Magnoliaceae. Our genome syntenic and Ks distribution analyses suggested a shared WGD event by *Magnolia* and *Liriodendron*. As the timing of this WGD is around ~116 MYA estimated by Chen *et al.*^12^ and ~124 MYA in our study, this WGD appears to be shared even by the two sister families of Magnoliaceae and Annonaceae as the two lineages diverged at around 95–113 MYA (mean, 104 MYA) according to Timetree web service (www.timetree.org) and 56.5–91.5 MYA (mean, ~72.8 MYA) in our dating analysis. However, the soursop (*Annona muricata*, Annonaceae) genome has only a small ambiguous Ks peak (possibly indicating a small-scale duplication event rather than WGD^10^) detected at around 1.3–1.5 Ks units, which is even older than the divergence of Magnoliales and Laurales at around 1.0–1.1 Ks units, thus rejecting the possibility of a Magnoliaceae and Annonaceae shared WGD^10^. As the estimated divergence of *Liriodendron* and *Magnolia*/*Annona* occurred at around 0.18 and 0.6–0.7 Ks units (near the Ks peak of 0.75 in our study)^10^, respectively, this Magnoliaceae specific WGD might have possibly happened shortly after the split of Magnoliaceae and Annonaceae. Further, cytological evidences also support this Magnoliaceae specific WGD event. Annonaceae have a basic chromosome number of n=7, which is reported to be the original chromosome number for Magnoliales^85^, whereas the base number of Magnoliaceae is n=19, suggesting probable paleopolyploidy origin of Magnoliaceae. It is also worth noting that WGD events do not necessarily generate more species diversity in Magnoliales as the putatively WGD-depauperate Annonaceae with some 2,100 species is the largest family in Magnoliales in contrast to Magnoliaceae with a confirmed lineage specific WGD event whereas holding only ~300 members.

As a medicinal plant, the major effective component of the flower buds of *M. biondii* is the volatile oils constituted by a rich array of terpenoids, mainly sesquiterpenoids and monoterpenoids^86^. TPS genes of subfamily TPS-a and TPS-b are mainly responsible for the biosynthesis of sesquiterpenoids and monoterpenoids in mesangiosperms, respectively. Gene tree topologies for three angiosperm TPS proteins and comparison of TPS subfamily members with that of the other angiosperms^11^ revealed expansion of TPS genes in *M. biondii*, especially TPS-a and TPS-b subfamilies. Expression profiles of TPS genes in different tissues identified 33 TPS genes, primarily of TPS-a and TPS-b subfamilies, substantially expressed in flowers, compared to leaves. The expansions and significant expressions of these TPS genes in the subfamilies TPS-a and TPS-b in *M. biondii* is in concert with the high accumulation of sesquiterpenoids and monoterpenoids in the volatile oils extracted from the flower buds of *M. biondii*^86^.

## Conclusion

We constructed a reference genome of *M. biondii* by combining 10X Genomics Chromium, single-molecule real-time sequencing (SMRT), and Hi-C scaffolding strategies. The ~2.22 Gb genome assembly of *M. biondii*, with a heterozygosity of 0.73% and a repeat ratio of 66.48%, represented the largest genome among six sequenced genomes of magnoliids. We predicted a total of 48,319 protein genes from the genome assembly of *M. biondii*, 81.55% of which were functionally annotated. Phylogenomic reconstruction strongly supported the sister relationship of magnoliids and Chloranthaceae, which together formed a sister relationship with a clade comprising monocots and eudicots. Our new genome information should further enhance the knowledge on the molecular basis of genetic diversity and individual traits in *Magnolia*, as well as the molecular breeding and early radiations of angiosperms.

## Acknowledgements

This work was supported the National Key R&D Program of China (No. 2019YFC1711000), the National Natural Science Foundation (No. 31600171), the Shenzhen Urban Management Bureau Fund (No. 201520), and the Shenzhen Municipal Government of China (No. JCYJ20170817145512467). This work is part of the 10KP project. We sincerely thank the support provided by China National GeneBank.

## Authors’ contributions

S.Z. and H.L. designed and coordinated the whole project. M.L., S.D., S.Z. and H.L. together led and performed the whole project. M.L, S.D., and F.C. performed the analyses of genome evolution, gene family analyses. S.D., M.L., H.L., S.Z., Y.L., X.G., and E.W. participated in the manuscript writing and revision. All authors read and approved the final manuscript.

## Supplementary Information

Supplementary Information accompanies this paper at XXX.

