## Supplementary figures for "The genome assembly and annotation of *Magnolia biondii* Pamp., a phylogenetically, economically, and medicinally important ornamental tree species"

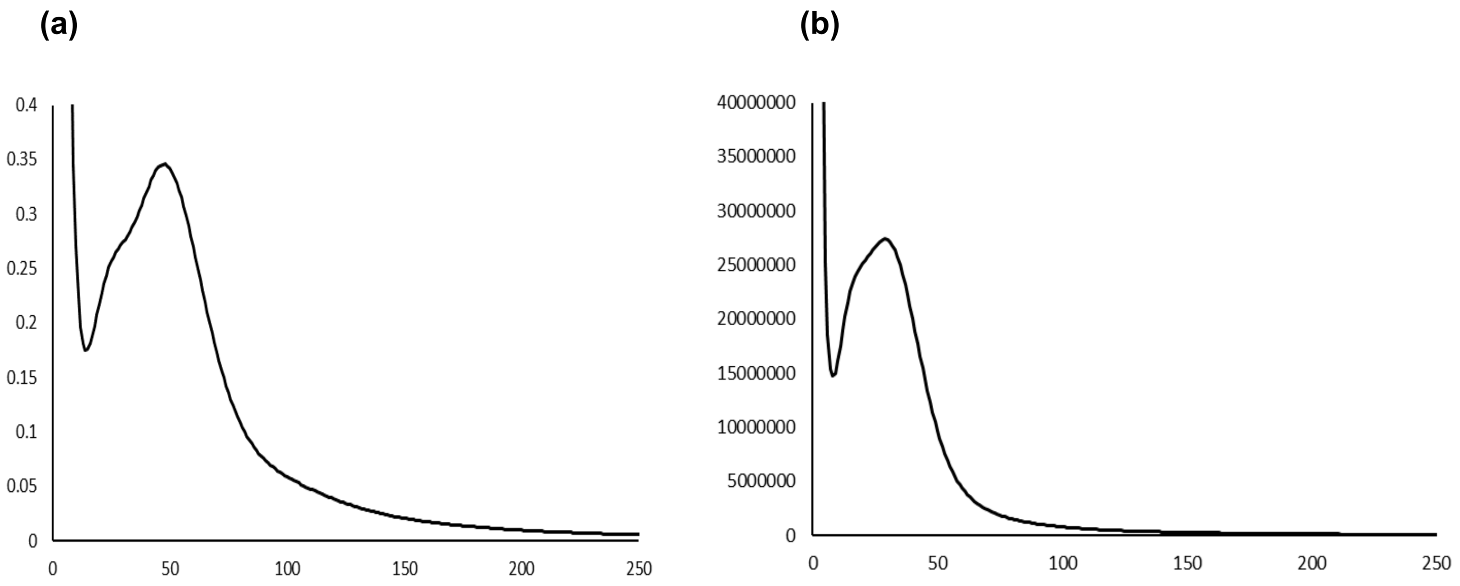


**Figure S1.** Estimation of the genome size of the *M. biondii* by (a) K-mer frequency analysis, and (b) GCE software

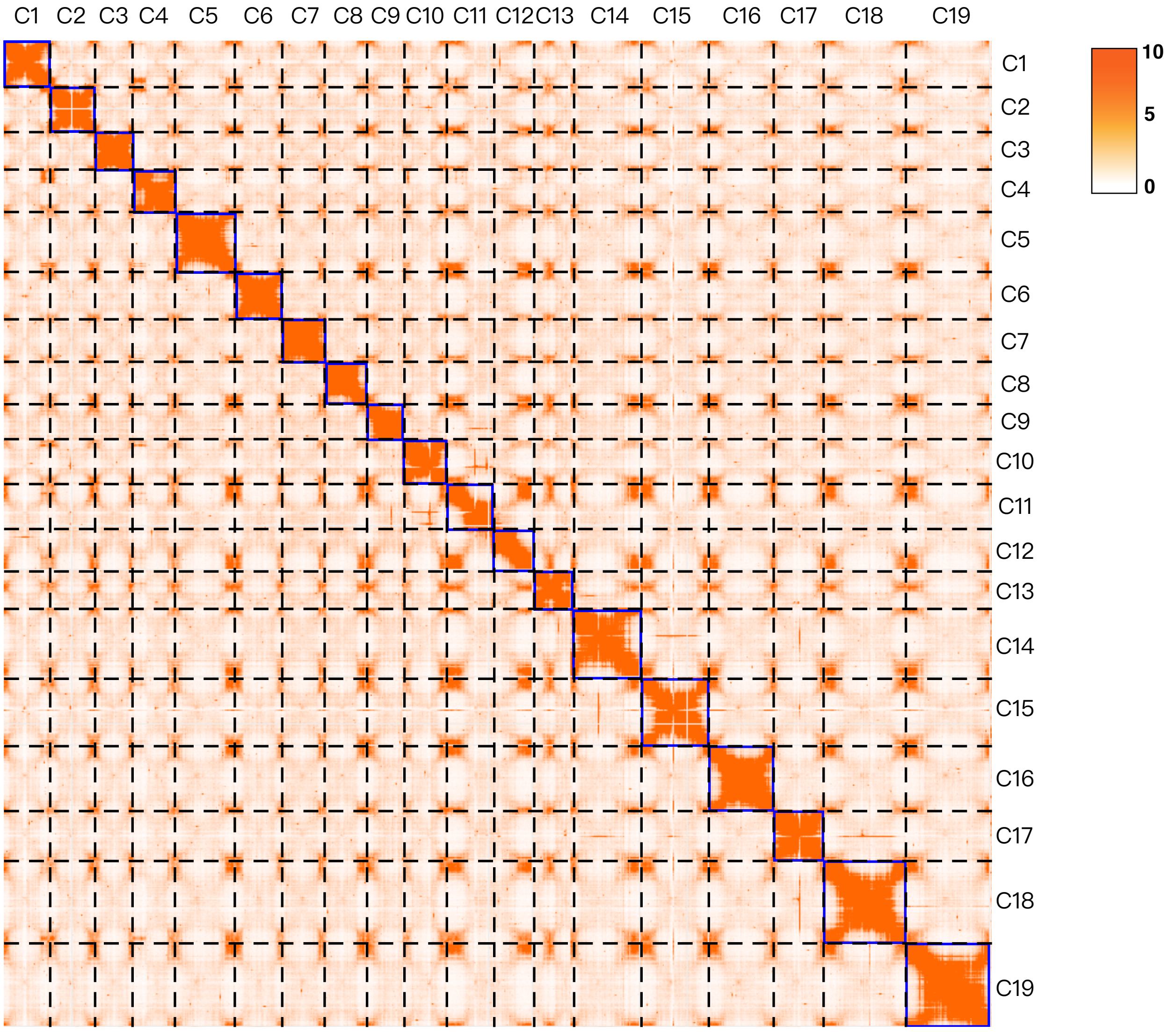


**Figure S2.** Hi-C map of the *M. biondii* genome showing genome-wide all-by-all interactions. The map shows a high resolution of individual chromosomes that are scaffolded and assembled independently. The heat map colors ranging from light yellow to dark red indicate the frequency of Hi-C interaction links from low to high (0–10).

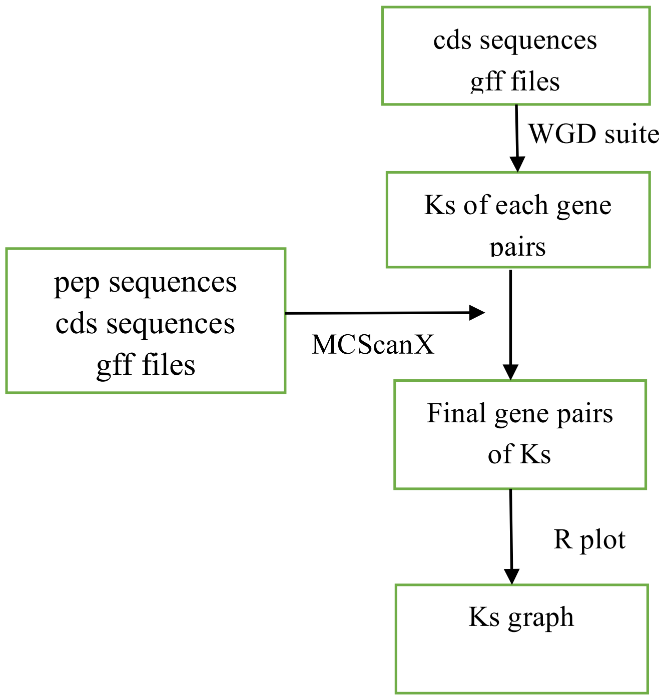


**Figure S3.** The flow chart displaying the analyses of WGD.

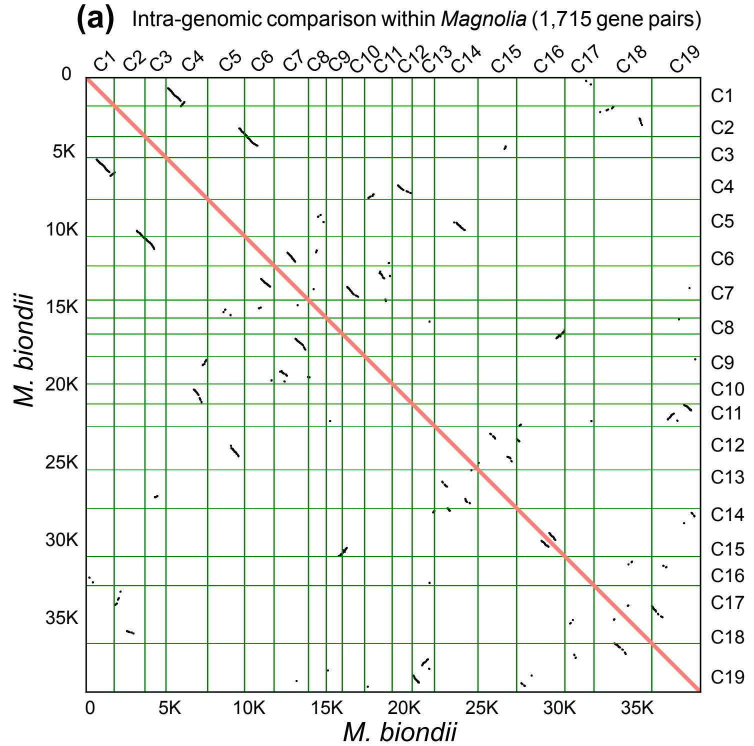


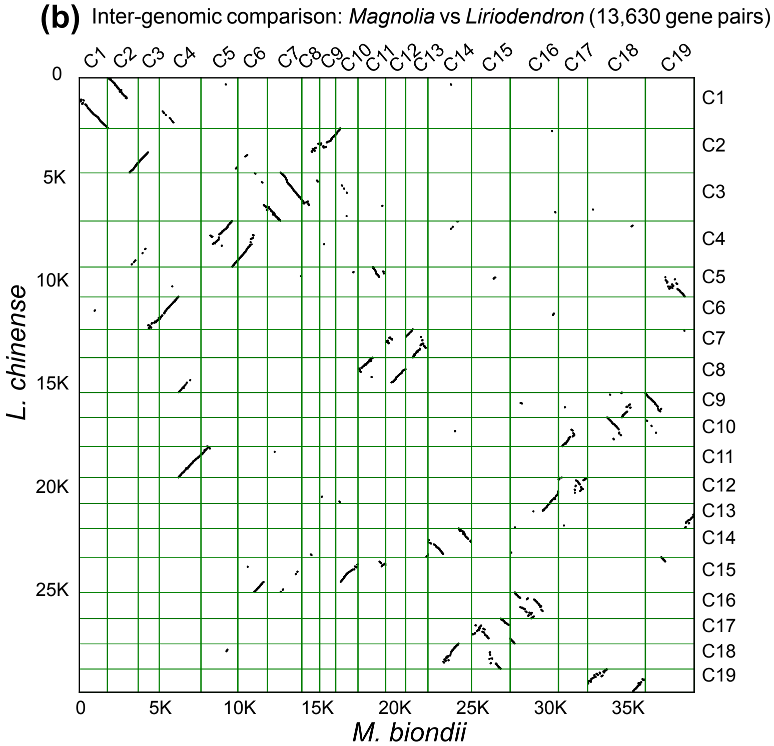


**Figure S4.** Intra-genomic comparison (a) within *M. biondii*. (b). of *M. biondii* and *L. chinense* genomes.
