## Supplementary tables for "The genome assembly and annotation of *Magnolia biondii* Pamp., a phylogenetically, economically, and medicinally important ornamental tree species"

**Table S1. Sequencing statistics.**

| DNA-seq | Raw data | | | | | Clean data | | | |
|---|---|---|---|---|---|---|---|---|---|
| | Library type | Read length (max; mean) | Total number of reads | Total number of bases (Gb) | Depth (X) | Read length (bp) | Total number of reads (bp) | Total number of bases (Gb) | Depth (X) |
| 10 x Genomics | Chromium | 150 | 1,169,695,614 | 175.45 | 80.00 | 150 | 908,810,764 | 136.32 | 61.96 |
| Pacific SMRT | 20kb library | 100,846;10324 | 5,953,700 | 66.78 | 30.35 | NA | NA | NA | NA |
| HI-C | | 100 | 768,917,688 | 153.78 | 69.9 | | | | |

| RNA-seq | Raw data | | Clean data | | Sample |
|---|---|---|---|---|---|
| | Read number (bp) | Base number (bp) | Base number (bp) | Reads number (bp) | |
| FLOWER | 51,130,512 | 4,601,746,080 | 2,898,235,224 | 33,897,488 | flower |
| BUDA | 51,847,396 | 4,666,265,640 | 3,154,682,214 | 36,896,868 | pre-meiosis flower |
| BUDB | 52,557,176 | 4,730,145,840 | 2,609,209,314 | 30,517,068 | post-meiosis flower |
| LEAF | 51,326,726 | 4,619,405,340 | 2,197,638,135 | 25,703,370 | leaf |
| Total | 206,861,810 | 18,617,562,900 | 10,859,764,887 | 127,014,794 | / |

**Table S2. K-mer survey statistics and evaluation of Hi-C data**

**K-mer frequency analysis**

| Kmer | Kmer_num | Pkdepth | Genome size (Mb) | Used base (Gb) | Used read (Mb) | X |
|---|---|---|---|---|---|---|
| 17 | 104,335,244,432 | 48 | 2,173.65 | 118.35 | 876.39 | 54.451 |

**GCE software**

| Raw_peak | Now_node | Cvg | Genome size (Mb) | Heterozygous ratio (%) | Repeat ratio (%) |
|---|---|---|---|---|---|
| 29 | 1,113,015,883 | 29.5297 | 2,236.75 | 0.73 | 61.83 |

**Evaluation of Hi-C reads**

| total reads pairs | 768917688 |
|---|---|
| reads1 mapping rate | 0.8556 |
| reads2 mapping rate | 0.8558 |
| valid reads pairs | 281232993 |
| Pair Type %(L-I-O-R) | 25% - 25% - 25% - 25% |
| unique reads | 171064287 |
| Hi-C contacts reads | 158069682 |
| Long Range (>20Kb) reads | 24336227 |

**Table S3. Statistics of the raw genome assemblies.**

| | Canu + corrected | wtdbg + raw reads | wtdbg + corrected reads | SamrtDenovo+ corrected | Flye | Miniasm |
|---|---|---|---|---|---|---|
| Total Contig length (Gb) | 1.87 | 1.75 | 1.53 | 1.48 | 1.78 | 2.20 |
| Number of contigs | 16,431 | 14,273 | 12,028 | 14,204 | 14,263 | 15,713 |
| Contig N50 (Kb) | 185.16 | 265.34 | 253.33 | 147.33 | 275.71 | 267.112 |
| Contig N90 (Kb) | 47.29 | 60.17 | 62.11 | 47.68 | 60.76 | 59.46 |
| Max contig length(Kb) | 1,848.39 | 2,202.97 | 1,532.87 | 1,049.52 | 2,205.34 | 2,125.10 |
| Complete BUSCOs (%) | 61.5 | 32.1 | 60.8 | 4.9 | 72.0 | 75.6 |
| Complete and single-copy BUSCOs (%) | 57.7 | 31.8 | 59.5 | 4.6 | 69.3 | 73.5 |
| Complete and duplicated BUSCOs (%) | 3.8 | 0.3 | 1.3 | 0.3 | 2.7 | 2.1 |
| Fragmented BUSCOs (%) | 16.0 | 22.1 | 18.5 | 10.8 | 12.9 | 10.0 |

**Table S4. Repeat annotations of the *M. biondii* genome assembly**

| Repeat elements | Type | % of genome | Length (bp) |
|---|---|---|---|
| Type I: Retrotransposon elements | SINE | 0.01 | 252,416 |
| | LINE | 4.47 | 99,544,077 |
| | LTR | 58.06 | 1,291,718,910 |
| | LTR/Copia | 26.28 | 584,683,554 |
| | LTR/Gypsy | 29.42 | 654,355,377 |
| Type II: DNA transposon | DNA | 5.86 | 130,503,028 |
| Type III: Tandem repeats | Satellite | 0.24 | 5,540,573 |
| | Simple_repeat | 0.79 | 17,626,796 |
| Others | | 0.32 | 7,240,517 |
| Total repeat | | 66.48 | 1,478,819,185 |

### Table S5. Gene annotation statistics of the *M. biondii* assembly and transcriptome assembly statistics

#### Protein-coding genes prediction

| Species | Genome size (Mb) | Gene number | Average /median gene length (bp) | Average /median coding sequence length (bp) | Average exons per gene | Average /median exon length (bp) | Average /median intron length (bp) | C (Complete BUSCOs) | S (Complete and single-copy BUSCOs) | D (Complete and duplicated BUSCOs) | F (Fragmented BUSCOs) | M (Missing BUSCOs) |
|---|---|---|---|---|---|---|---|---|---|---|---|---|
| ***M. biondii*** | **2,224** | **48,319** | **10,576/3,390** | **950/669** | **4.44** | **214/132** | **2,797/532** | **1,232 (89.60%)** | **1,171 (85.16%)** | **61 (4.44%)** | **78 (5.67%)** | **65 (4.73%)** |
| *A. trichopoda* | 706 | 31,058 | 12,633/7,461 | 1,457/1,206 | 6.96 | 209/118 | 1,876/487 | 1,351 (98.25%) | 672 (48.87%) | 679 (49.38%) | 9 (0.65%) | 15 (1.09%) |
| *A. thaliana* | 119 | 26,633 | 1,909/1,593 | 1,242/1,065 | 5.23 | 237/134 | 158/98 | 1,370 (99.63%) | 1,360 (98.91%) | 10 (0.73%) | 2 (0.14%) | 3 (0.22%) |
| *C. kanehirae* | 730 | 26,512 | 7,609/4,610 | 1,320/1,095 | 5.42 | 244/135 | 1,422/524 | 1,242 (90.33%) | 1,160 (84.36%) | 82 (5.96%) | 39 (2.84%) | 94 (6.84%) |
| *L. chinense* | 1,742 | 30,787 | 10,740/5,792 | 1,271/1,080 | 4.93 | 258/140 | 2,411/587 | 1,109 (80.65%) | 1,016 (73.89%) | 93 (6.76%) | 164 (11.93%) | 102 (7.42%) |
| *O. sativa* | 374 | 40,852 | 3,418/2,547 | 1,421/1,212 | 5.67 | 251/128 | 428/145 | 1,371 (99.71%) | 1,020 (74.18%) | 351 (25.53%) | 0 (0%) | 4 (0.29%) |

#### Non-coding RNA genes in the genome of *M. biondii*

| | | | rRNA | | | | | snRNA | | | |
|---|---|---|---|---|---|---|---|---|---|---|---|
| Type | miRNA | tRNA | rRNA | 18S | 28S | 5.8S | 5S | snRNA | CD-box | HACA-box | splicing |
| Copy (w) | 109 | 904 | 959 | 243 | 480 | 98 | 138 | 3,713 | 3,383 | 67 | 263 |
| Average length (bp) | 124 | 75 | 292 | 765 | 134 | 153 | 104 | 110 | 106 | 134 | 153 |
| Total length (bp) | 13,493 | 68,172 | 279,809 | 185,909 | 64,541 | 15,019 | 14,340 | 407,934 | 358,695 | 8,973 | 40,266 |
| % of genome | 0.000607 | 0.003065 | 0.012578 | 0.008357 | 0.002901 | 0.000675 | 0.000645 | 0.018338 | 0.016125 | 0.000403 | 0.001810 |

### Transcriptome assembly

| | FLOWER | BUDA | BUDB | LEAF |
|---|---|---|---|---|
| Complete BUSCOs | 719 (52.29%) | 880 (64.00%) | 847 (61.60%) | 753 (54.76%) |
| Complete and Single-copy BUSCOs | 563 (40.95%) | 672 (48.87%) | 612 (44.51%) | 583 (42.40%) |
| Complete and Duplicated BUSCOs | 156 (11.35%) | 208 (15.13%) | 235 (17.09%) | 170 (12.36%) |
| Fragmented BUSCOs | 313 (22.76%) | 300 (21.82%) | 339 (24.65%) | 354 (25.75%) |
| Missing BUSCOs | 343 (24.95%) | 195 (14.18%) | 189 (13.75%) | 268 (19.49%) |

**Table S6. Functional annotation of predicted genes in *M. biondii* genome**

| Values | NR | Swissprot | KEGG | COG | TrEMBL | Interpro | GO | Overall | Unannotated |
|---|---|---|---|---|---|---|---|---|---|
| Number | 38,784 | 28,197 | 28,545 | 12,602 | 38,050 | 28,639 | 17,908 | 39,405 | 8,914 |
| Percentage | 80.27% | 58.36% | 59.08% | 26.08% | 78.75% | 59.27% | 37.06% | 81.55% | 18.45% |

**Table S7. Data used for phylogenetic reconstruction of angiosperms.**

| Taxa | Database | Url |
|---|---|---|
| *Amborella trichopoda* | AMTR1.0 | http://plants.ensembl.org/Amborella_trichopoda/Info/Index |
| *Arabidopsis thaliana* | TAIR10 | TAIR10 |
| *Cinnamomum kanehirae* | GCA_003546025.1 | https://www.ncbi.nlm.nih.gov/genome/57158?genome_assembly_id=405569 |
| *Liriodendron chinense* | | http://120.78.193.56:8000/node/141281 |
| *Oryza sativa* | TIGR rice v4.0 | http://rice.plantbiology.msu.edu/ |
| *Picea abies* | Picea abies v1.0 | ftp://plantgenie.org/Data/ConGenIE/ |
| *Sorghum bicolor* | GCF 000003195.3 | https://www.ncbi.nlm.nih.gov/assembly/GCF_000003195.3 |
| *Vitis vinifera* | 12X v0 | http://www.genoscope.cns.fr/externe/Download/Projets/Projet_ML/data/12X/ |
| *Annona muricata* | OneKp | https://sites.google.com/a/ualberta.ca/onekp/ |
| *Aristolochia elegans* | OneKp | https://sites.google.com/a/ualberta.ca/onekp/ |
| *Ascarina rubricaulis* | OneKp | https://sites.google.com/a/ualberta.ca/onekp/ |
| *Canella winterana* | OneKp | https://sites.google.com/a/ualberta.ca/onekp/ |
| *Illicium floridanum* | OneKp | https://sites.google.com/a/ualberta.ca/onekp/ |
| *Illicium parviflorum* | OneKp | https://sites.google.com/a/ualberta.ca/onekp/ |
| *Laurelia sempervirens* | OneKp | https://sites.google.com/a/ualberta.ca/onekp/ |
| *Magnolia grandiflora* | OneKp | https://sites.google.com/a/ualberta.ca/onekp/ |
| *Nymphaea sp* | OneKp | https://sites.google.com/a/ualberta.ca/onekp/ |
| *Sarcandra glabra* | OneKp | https://sites.google.com/a/ualberta.ca/onekp/ |
| *Saruma henryi* | OneKp | https://sites.google.com/a/ualberta.ca/onekp/ |

**Table S8. KEGG enrichment of *M. biondii* unique gene families**

| Pathway ID | Pathway | Gene number | Total gene | P-value | negLog10_P value |
|---|---|---|---|---|---|
| ko03040 | Spliceosome | 669 | 1,342 | 5.16E-184 | 183.287647 |
| ko03030 | DNA replication | 625 | 1,213 | 7.05E-181 | 180.151837 |
| ko04120 | Ubiquitin mediated proteolysis | 439 | 1,005 | 6.69E-94 | 93.1747474 |
| ko03018 | RNA degradation | 386 | 842 | 5.85E-90 | 89.2327661 |
| ko00460 | Cyanoamino acid metabolism | 273 | 484 | 2.14E-89 | 88.6694965 |
| ko00940 | Phenylpropanoid biosynthesis | 294 | 807 | 4.23E-43 | 42.3740438 |
| ko03013 | RNA transport | 287 | 841 | 4.43E-36 | 35.3537527 |
| ko00500 | Starch and sucrose metabolism | 323 | 1,025 | 5.84E-33 | 32.2336403 |
| ko04626 | Plant-pathogen interaction | 486 | 1,808 | 5.91E-30 | 29.2286105 |
| ko00960 | Tropane, piperidine and pyridine alkaloid biosynthesis | 74 | 131 | 4.10E-25 | 24.3874105 |
| ko00130 | Ubiquinone and other terpenoid-quinone biosynthesis | 72 | 150 | 4.82E-19 | 18.3170777 |
| ko00950 | Isoquinoline alkaloid biosynthesis | 76 | 164 | 6.83E-19 | 18.1654283 |
| ko00400 | Phenylalanine, tyrosine and tryptophan biosynthesis | 82 | 186 | 1.32E-18 | 17.8806953 |
| ko01110 | Biosynthesis of secondary metabolites | 599 | 2,720 | 1.38E-14 | 13.8596378 |
| ko00350 | Tyrosine metabolism | 78 | 201 | 4.63E-14 | 13.3341398 |
| ko00270 | Cysteine and methionine metabolism | 82 | 242 | 4.45E-11 | 10.3513947 |
| ko00591 | Linoleic acid metabolism | 61 | 164 | 1.97E-10 | 9.70584296 |
| ko00730 | Thiamine metabolism | 23 | 41 | 1.08E-08 | 7.96600681 |
| ko00062 | Fatty acid elongation | 40 | 105 | 1.15E-07 | 6.93825884 |
| ko00902 | Monoterpenoid biosynthesis | 28 | 65 | 4.79E-07 | 6.31987189 |
| ko03410 | Base excision repair | 70 | 238 | 6.45E-07 | 6.19059215 |
| ko00360 | Phenylalanine metabolism | 81 | 289 | 8.01E-07 | 6.09629316 |
| ko04144 | Endocytosis | 111 | 445 | 4.72E-06 | 5.32637915 |
| ko03420 | Nucleotide excision repair | 54 | 193 | 5.32E-05 | 4.27371644 |
| ko00640 | Propanoate metabolism | 51 | 199 | 0.000831 | 3.08040838 |
| ko00280 | Valine, leucine and isoleucine degradation | 55 | 235 | 0.004694 | 2.3284756 |
| ko00410 | beta-Alanine metabolism | 45 | 192 | 0.009521 | 2.02130156 |
| ko03440 | Homologous recombination | 27 | 107 | 0.015345 | 1.81403056 |

**Table S9. GO enrichment of *M. biondii* unique gene families.**

| GO ID | GO Term | GO Class | P-value | Adjusted Pvalue | Gene number | Total gene |
|---|---|---|---|---|---|---|
| GO:0003676 | nucleic acid binding | MF | 9.87E-315 | 7.27E-312 | 1,007 | 3,397 |
| GO:0004523 | ribonuclease H activity | MF | 2.07E-188 | 7.63E-186 | 285 | 584 |
| GO:0004521 | endoribonuclease activity | MF | 1.60E-176 | 2.36E-174 | 287 | 616 |
| GO:0004540 | ribonuclease activity | MF | 2.02E-175 | 2.48E-173 | 288 | 622 |
| GO:0004518 | nuclease activity | MF | 3.48E-154 | 3.21E-152 | 292 | 689 |
| GO:0097159 | organic cyclic compound binding | MF | 1.55E-118 | 1.15E-116 | 1,230 | 6,768 |
| GO:1901363 | heterocyclic compound binding | MF | 1.56E-118 | 1.15E-116 | 1,230 | 6,768 |
| GO:0016788 | hydrolase activity, acting on ester bonds | MF | 1.17E-99 | 7.21E-98 | 305 | 941 |
| GO:0005488 | binding | MF | 1.20E-55 | 6.31E-54 | 1,516 | 10,651 |
| GO:0008725 | DNA-3-methyladenine glycosylase activity | MF | 3.12E-13 | 4.99E-12 | 20 | 29 |
| GO:0016787 | hydrolase activity | MF | 2.51E-11 | 2.31E-10 | 410 | 2,756 |
| GO:0046983 | protein dimerization activity | MF | 2.01E-05 | 9.44E-05 | 68 | 374 |
| GO:0003677 | DNA binding | MF | 0.000216996 | 0.00094074 | 168 | 1,156 |
| GO:0015930 | glutamate synthase activity | MF | 0.00261925 | 0.009798918 | 6 | 14 |
| GO:0016742 | hydroxymethyl-, formyl- and related transferase activity | MF | 0.003956081 | 0.014292312 | 6 | 15 |
| GO:0045735 | nutrient reservoir activity | MF | 0.006038772 | 0.020700347 | 17 | 79 |
| GO:0004602 | glutathione peroxidase activity | MF | 0.006987864 | 0.023732975 | 5 | 12 |
| GO:0016838 | carbon-oxygen lyase activity, acting on phosphates | MF | 0.007450789 | 0.024960143 | 21 | 106 |
| GO:0010333 | terpene synthase activity | MF | 0.010668804 | 0.034186559 | 20 | 102 |
| GO:0003871 | 5-methyltetrahydropteroyltriglutamate-homocysteine S-methyltransferase activity | MF | 0.014602509 | 0.04392673 | 5 | 14 |
| GO:0005337 | nucleoside transmembrane transporter activity | MF | 0.014602509 | 0.04392673 | 5 | 14 |
| GO:0006260 | DNA replication | BP | 4.24E-11 | 3.72E-10 | 30 | 89 |
| GO:0006284 | base-excision repair | BP | 2.03E-10 | 1.55E-09 | 20 | 37 |
| GO:0006259 | DNA metabolic process | BP | 1.30E-07 | 7.72E-07 | 61 | 290 |
| GO:0042147 | retrograde transport, endosome to Golgi | BP | 0.001482682 | 0.005751244 | 5 | 9 |
| GO:0006904 | vesicle docking involved in exocytosis | BP | 0.002244019 | 0.008481243 | 7 | 18 |
| GO:0006281 | DNA repair | BP | 0.003109873 | 0.011402868 | 25 | 126 |
| GO:0000786 | nucleosome | CC | 1.07E-07 | 6.44E-07 | 26 | 88 |
| GO:0044427 | chromosomal part | CC | 1.45E-07 | 8.47E-07 | 32 | 120 |
| GO:0005694 | chromosome | CC | 8.68E-06 | 4.41E-05 | 33 | 142 |
| GO:0005664 | nuclear origin of replication recognition complex | CC | 0.000542268 | 0.002296848 | 6 | 11 |
| GO:0030906 | retromer complex, inner shell | CC | 0.001482682 | 0.005751244 | 5 | 9 |

**Table S10. KEGG enrichment of the expanded gene families.**

| Pathway ID | Pathway | Gene number | Total gene | P-value |
|---|---|---|---|---|
| ko02010 | ABC transporters | 211 | 286 | 2.51E-32 |
| ko00052 | Galactose metabolism | 191 | 277 | 1.80E-23 |
| ko00945 | Stilbenoid, diarylheptanoid and gingerol biosynthesis | 99 | 118 | 2.76E-23 |
| ko00941 | Flavonoid biosynthesis | 135 | 193 | 7.55E-18 |
| ko00604 | Glycosphingolipid biosynthesis - ganglio series | 64 | 74 | 6.65E-17 |
| ko00531 | Glycosaminoglycan degradation | 78 | 98 | 4.86E-16 |
| ko04710 | Circadian rhythm - mammal | 6 | 78 | 1.68E-11 |
| ko00600 | Sphingolipid metabolism | 79 | 116 | 3.76E-10 |
| ko03020 | RNA polymerase | 118 | 194 | 1.30E-09 |
| ko03008 | Ribosome biogenesis in eukaryotes | 188 | 344 | 6.84E-09 |
| ko00561 | Glycerolipid metabolism | 117 | 198 | 1.65E-08 |
| ko00511 | Other glycan degradation | 89 | 142 | 1.76E-08 |
| ko00510 | N-Glycan biosynthesis | 97 | 161 | 7.18E-08 |
| ko00904 | Diterpenoid biosynthesis | 48 | 68 | 1.90E-07 |
| ko00240 | Pyrimidine metabolism | 212 | 421 | 3.10E-06 |
| ko00040 | Pentose and glucuronate interconversions | 131 | 244 | 4.25E-06 |
| ko03050 | Proteasome | 115 | 213 | 1.13E-05 |
| ko04075 | Plant hormone signal transduction | 349 | 742 | 1.23E-05 |
| ko00402 | Benzoxazinoid biosynthesis | 16 | 18 | 2.06E-05 |
| ko00906 | Carotenoid biosynthesis | 67 | 114 | 2.26E-05 |
| ko03010 | Ribosome | 363 | 779 | 2.27E-05 |
| ko00944 | Flavone and flavonol biosynthesis | 39 | 60 | 5.44E-05 |
| ko00053 | Ascorbate and aldarate metabolism | 49 | 81 | 0.00010085 |
| ko01100 | Metabolic pathways | 2,020 | 4,848 | 0.00026593 |
| ko00603 | Glycosphingolipid biosynthesis - globo series | 21 | 29 | 0.00032564 |
| ko00592 | alpha-Linolenic acid metabolism | 65 | 119 | 0.00055365 |
| ko00190 | Oxidative phosphorylation | 141 | 288 | 0.00061416 |
| ko00900 | Terpenoid backbone biosynthesis | 61 | 112 | 0.00088856 |
| ko00195 | Photosynthesis | 75 | 144 | 0.00137519 |
| ko00902 | Monoterpenoid biosynthesis | 37 | 65 | 0.00320013 |
| ko00901 | Indole alkaloid biosynthesis | 12 | 16 | 0.00425485 |
| ko01040 | Biosynthesis of unsaturated fatty acids | 45 | 86 | 0.01029769 |
| ko00450 | Selenocompound metabolism | 42 | 80 | 0.01201263 |
| ko03015 | mRNA surveillance pathway | 161 | 361 | 0.02520815 |
| ko00480 | Glutathione metabolism | 82 | 175 | 0.0270918 |
| ko00520 | Amino sugar and nucleotide sugar metabolism | 99 | 215 | 0.02806702 |
| ko00966 | Glucosinolate biosynthesis | 14 | 23 | 0.0307158 |
| ko00908 | Zeatin biosynthesis | 41 | 83 | 0.04136432 |
| ko00073 | Cutin, suberine and wax biosynthesis | 39 | 79 | 0.04628199 |

**Table S11. GO (level 3) enrichment of the expanded gene families.**

| GO ID | GO Term | GO Class | Pvalue | Adjusted Pvalue | Gene number | Total gene |
|---|---|---|---|---|---|---|
| GO:0016491 | oxidoreductase activity | MF | 6.34E-39 | 9.38E-37 | 908 | 1,521 |
| GO:0016740 | transferase activity | MF | 8.46E-18 | 3.13E-16 | 1,436 | 2,805 |
| GO:0004601 | peroxidase activity | MF | 2.08E-15 | 6.14E-14 | 107 | 138 |
| GO:0022857 | transmembrane transporter activity | MF | 1.01E-14 | 2.49E-13 | 402 | 691 |
| GO:0016829 | lyase activity | MF | 9.81E-14 | 1.61E-12 | 174 | 261 |
| GO:0043167 | ion binding | MF | 2.38E-12 | 2.94E-11 | 2,403 | 5,010 |
| GO:0090484 | drug transporter activity | MF | 6.93E-11 | 7.33E-10 | 55 | 65 |
| GO:0022892 | substrate-specific transporter activity | MF | 8.38E-08 | 5.17E-07 | 177 | 299 |
| GO:0030246 | carbohydrate binding | MF | 4.46E-06 | 2.44E-05 | 95 | 152 |
| GO:0045156 | electron transporter, transferring electrons within the cyclic electron transport pathway of photosynthesis activity | MF | 0.000258091 | 0.00119367 | 10 | 10 |
| GO:0004857 | enzyme inhibitor activity | MF | 0.000342449 | 0.001490662 | 62 | 100 |
| GO:0097367 | carbohydrate derivative binding | MF | 0.002358666 | 0.009186384 | 16 | 20 |
| GO:0038023 | signaling receptor activity | MF | 0.002860228 | 0.010854199 | 35 | 54 |
| GO:0004871 | signal transducer activity | MF | 0.004621154 | 0.016681238 | 46 | 76 |
| GO:0042221 | response to chemical stimulus | BP | 1.95E-24 | 1.44E-22 | 145 | 177 |
| GO:0044710 | single-organism metabolic process | BP | 7.52E-19 | 3.71E-17 | 1,117 | 2,116 |
| GO:0044765 | single-organism transport | BP | 4.57E-14 | 9.66E-13 | 638 | 1,173 |
| GO:0009719 | response to endogenous stimulus | BP | 9.70E-14 | 1.61E-12 | 78 | 95 |
| GO:0044707 | single-multicellular organism process | BP | 2.36E-10 | 2.32E-09 | 78 | 104 |
| GO:0051234 | establishment of localization | BP | 2.63E-10 | 2.43E-09 | 728 | 1,404 |
| GO:0022414 | reproductive process | BP | 3.85E-09 | 3.35E-08 | 64 | 84 |
| GO:0044703 | multi-organism reproductive process | BP | 6.72E-09 | 5.23E-08 | 60 | 78 |
| GO:0044706 | multi-multicellular organism process | BP | 6.72E-09 | 5.23E-08 | 60 | 78 |
| GO:0071554 | cell wall organization or biogenesis | BP | 1.81E-08 | 1.28E-07 | 79 | 112 |
| GO:0048610 | cellular process involved in reproduction | BP | 3.25E-08 | 2.19E-07 | 60 | 80 |
| GO:0044763 | single-organism cellular process | BP | 6.03E-07 | 3.43E-06 | 1,033 | 2,114 |
| GO:0009607 | response to biotic stimulus | BP | 5.80E-06 | 3.07E-05 | 24 | 27 |
| GO:0016043 | cellular component organization | BP | 0.001089746 | 0.004480066 | 178 | 338 |
| GO:0007275 | multicellular organismal development | BP | 0.003994044 | 0.014777962 | 18 | 24 |
| GO:0051707 | response to other organism | BP | 0.013010792 | 0.043763573 | 8 | 9 |
| GO:0044238 | primary metabolic process | BP | 0.014402167 | 0.046745228 | 2,469 | 5,467 |
| GO:0031224 | intrinsic to membrane | CC | 3.20E-12 | 3.64E-11 | 474 | 856 |
| GO:0071944 | cell periphery | CC | 2.20E-07 | 1.30E-06 | 104 | 162 |
| GO:0030312 | external encapsulating structure | CC | 1.33E-05 | 6.80E-05 | 62 | 93 |
| GO:0044425 | membrane part | CC | 0.000101562 | 0.000484875 | 519 | 1,046 |
